# Harmonic imaging for nonlinear detection of acoustic biomolecules

**DOI:** 10.1101/2024.06.18.599141

**Authors:** Rohit Nayak, Mengtong Duan, Bill Ling, Zhiyang Jin, Dina Malounda, Mikhail G. Shapiro

## Abstract

Gas vesicles (GVs) based on acoustic reporter genes have emerged as potent contrast agents for cellular and molecular ultrasound imaging. These air-filled, genetically encoded protein nanostructures can be expressed in a variety of cell types *in vivo* to visualize cell location and activity or injected systemically to label and monitor tissue function. Distinguishing GVs from tissue signal deep inside intact organisms requires imaging approaches such as amplitude modulation (AM) or collapse-based pulse sequences, however they have limitations in sensitivity or require irreversible collapse of the GVs that restricts its scope for imaging dynamic cellular processes. To address these limitations, this study explores the utility of harmonic imaging to enhance the sensitivity of non-destructive imaging of GVs and cellular processes. Traditional fundamental-frequency imaging utilizing cross-wave AM (xAM) sequences has been deemed optimal for GV imaging. Contrary to this, we hypothesize that harmonic imaging, integrated with xAM could significantly elevate GV detection sensitivity. To verify our hypothesis, we conducted imaging on tissue-mimicking phantoms embedded with purified GVs, mammalian cells genetically modified to express GVs, and live mice after systemic GV infusion. Our findings reveal that harmonic xAM (HxAM) imaging markedly surpasses traditional xAM in isolating GVs’ nonlinear acoustic signature, showcasing significant enhancements in signal-to-background and contrast-to-background ratios across all tested samples. Further investigation into the backscattered spectra elucidates the efficacy of harmonic imaging in conjunction with xAM. HxAM imaging enables the detection of lower concentrations of GVs and cells with ultrasound and extends the imaging depth *in vivo* by up to 20% and imaging performance metrics by up to 10dB. These advancements bolster the capabilities of ultrasound for molecular and cellular imaging, underscoring the potential of using harmonic signals to amplify GV detection.

---

Ultrasound imaging plays a pivotal role in medical diagnostics, offering high spatial and temporal resolution for examining organ anatomy and function. The development of micro- and nanoscale contrast agents has extended ultrasound’s reach considerably^1^, while the introduction of acoustic reporter genes has broadened the capabilities of ultrasound imaging at the molecular and cellular levels^2^. In particular, gas vesicles (GVs), a naturally derived class of genetically encoded protein shells enclosing a compartment of air with dimensions of ∼100 nm, can serve as both purified nanoscale contrast agents and genetically encoded reporter genes and biosensors^3–10^. This enables their use across wide range of applications such as targeted cellular imaging and real-time monitoring in various biological contexts.

GVs are distinctive in their structure: typically, 100 nm in diameter and 500 nm in length, they are encapsulated by a protein shell about 3 nm thick that can endure large pressures up to hundreds of kilopascals, without collapsing^8,9^. The shell’s interior is markedly hydrophobic, preventing water ingress while permitting gas molecules to freely diffuse in and out^6,7^. The architectural integrity of GVs is maintained by a primary structural protein, GvpA, a small (8-kDa) and highly hydrophobic molecule that polymerizes to form the shell^6^. This protein assembly is further reinforced by a secondary protein, GvpC, which externally fortifies the shell, enhancing the GVs’ mechanical strength^10^. The low density and high compressibility of GVs enable effective sound wave scattering, generating substantial ultrasound backscatter^2^. This feature becomes crucial when GVs are heterologously expressed as ARGs in genetically engineered cells^3,4^, broadening their application from targeted cellular imaging to real-time monitoring of biological processes^11–15^. Distinguishing GVs backscatter from the tissue necessitates a method that separates these signals. The distinct mechanical behavior of GVs, marked by reversible nonlinear buckling beyond specific pressure thresholds, facilitates the use of amplitude modulation (AM) techniques for isolating their signals from background tissue clutter^16–18^. AM pulse sequences typically involve three sequential transmit pulses, with one at full amplitude and the others at half amplitude. The full-amplitude pulse is designed to elicit nonlinear behavior in GVs, which is minimal in tissue due to its substantially linear behavior. Subsequently, the half amplitude pulses elicit a linear response from both GVs and tissue. By subtracting the latter two transmissions from the first, tissue clutter is minimized, whereas the nonlinear response of the GVs is amplified. This process effectively enhances the visualization of cells expressing GVs or systemically injected GVs.

AM based on cross-propagating waves (xAM) has proven especially effective in detecting GVs while minimizing artifacts arising from nonlinear wave propagation through GV inclusions^17^. As an alternative, BURST imaging provides the most sensitive detection of GV by capturing the unique signals produced by their intentional acoustic collapse – enhancing detectability by an order of magnitude compared to AM^5,19^. However, the applicability of BURST is limited in contexts requiring preservation of the GVs, such as dynamic imaging or biosensing. Therefore, enhancing the detection sensitivity of AM-based imaging for GVs is a critical goal in biomolecular ultrasound.

Harmonic imaging has been a key approach in improving the contrast and resolution of ultrasound images. For example, tissue harmonic imaging is routinely used in diagnostic ultrasonography for generating images with superior tissue definition, improved signal-to-noise ratio and reduced artifacts produced by side lobes, grating lobes, and reverberation^20,21^. The principle behind harmonic imaging lies in its use of the higher-frequency harmonics generated by the nonlinear propagation of the fundamental ultrasound wave. These harmonic waves typically contain fewer artifacts compared to images produced using conventional fundamental wave ultrasound. Building on this approach, the integration of harmonic imaging with specialized pulse sequences such as AM and pulse inversion has enhanced the detection of ultrasound contrast agents^22^. The relatively strong harmonic signals arising from the agents’ nonlinear acoustic behavior help set them apart from surrounding tissue^23^. In fields such as cardiology and hepatic imaging, harmonic imaging provides refined views of myocardial perfusion and endocardial borders, as well as more accurate lesion characterization^20^.

In this study, we evaluate the potential of using harmonic signals to enhance the detection of GVs. While existing research suggests that GVs can produce harmonic scattering^2,11,24^, this observation has not been integrated with AM, and previous work suggested that fundamental-frequency imaging provided the best GV imaging performance with conventional parabolic AM (pAM) sequences^16^. We hypothesized that integrating harmonic imaging with xAM could significantly improve GV detection sensitivity due to the cleaner nonlinear background of xAM. To test our hypothesis, we imaged tissue-mimicking phantoms containing purified GVs^8^, mammalian cells genetically engineered to express GVs as acoustic reporter genes^5^, and live mice following systemic infusion and liver uptake of GVs^25^. Further, we compared the performance of xAM and its harmonic counterpart, throughout the study.

For all experiments, we employed a 128-element linear array ultrasound probe with a nominal bandwidth of 14-22 MHz. To integrate harmonic signals with xAM imaging, we performed tests at a transmit frequency of 12.5 MHz, where we expected to observe second harmonic contributions manifesting at 25 MHz [Fig. 1]. To evaluate the effectiveness of this approach, we compared the outcomes with those obtained from conventional xAM imaging, which was performed at a baseline transmit frequency of 15.625 MHz (whose second harmonic is beyond the bandwidth of the transducer). We ensured that both the 12.5 MHz and 15.625 MHz frequencies were operated at the same transmit pressure of approximately 400 kPa, facilitating a direct and consistent comparison between the two imaging modes.

**Figure 1:**
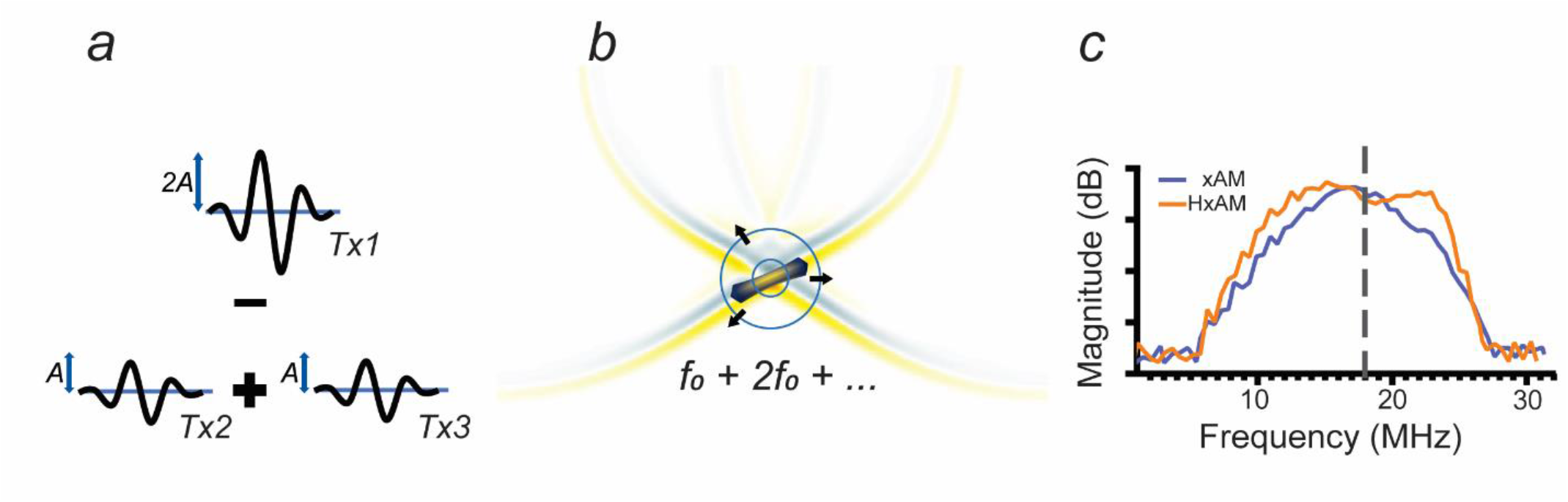
Cross-wave propagation and amplitude modulation in harmonic imaging. **(a)** Depicts a three-pulse sequence utilized for amplitude modulation, where the first pulse has twice the amplitude of the subsequent pulses to induce modulation. **(b)** Shows the nonlinear scattering behavior of harmonic gas GVs when sonicated above their buckling threshold, as indicated by the converging cross-waves. **(c)** Compares the receive spectra for ultrasound transmissions at 15.625 MHz and 12.5 MHz; the latter captures the harmonic signal within the bandwidth limitations of the transducer and scanner.

We began by imaging purified *Anabaena flos-aquae* GVs, stripped of GvpC to enable buckling, in tissue-mimicking phantoms^8^ [Fig. 2 (a)]. We conducted imaging of the samples using xAM, harmonic xAM (HxAM) using the full received signal, and a variant of harmonic xAM (HxAM-f) employing a high-pass receive filter set at 17.5 MHz to exclude the fundamental signal. We assessed the efficacy of imaging techniques at varying concentrations of GVs, determined by optical density (OD) measurements, which is based on light scattering. An OD of 1, measured at a wavelength of 500 nm (OD_500_), corresponds to a concentration of approximately 184 pM GV particles^8^.

**Figure 2:**
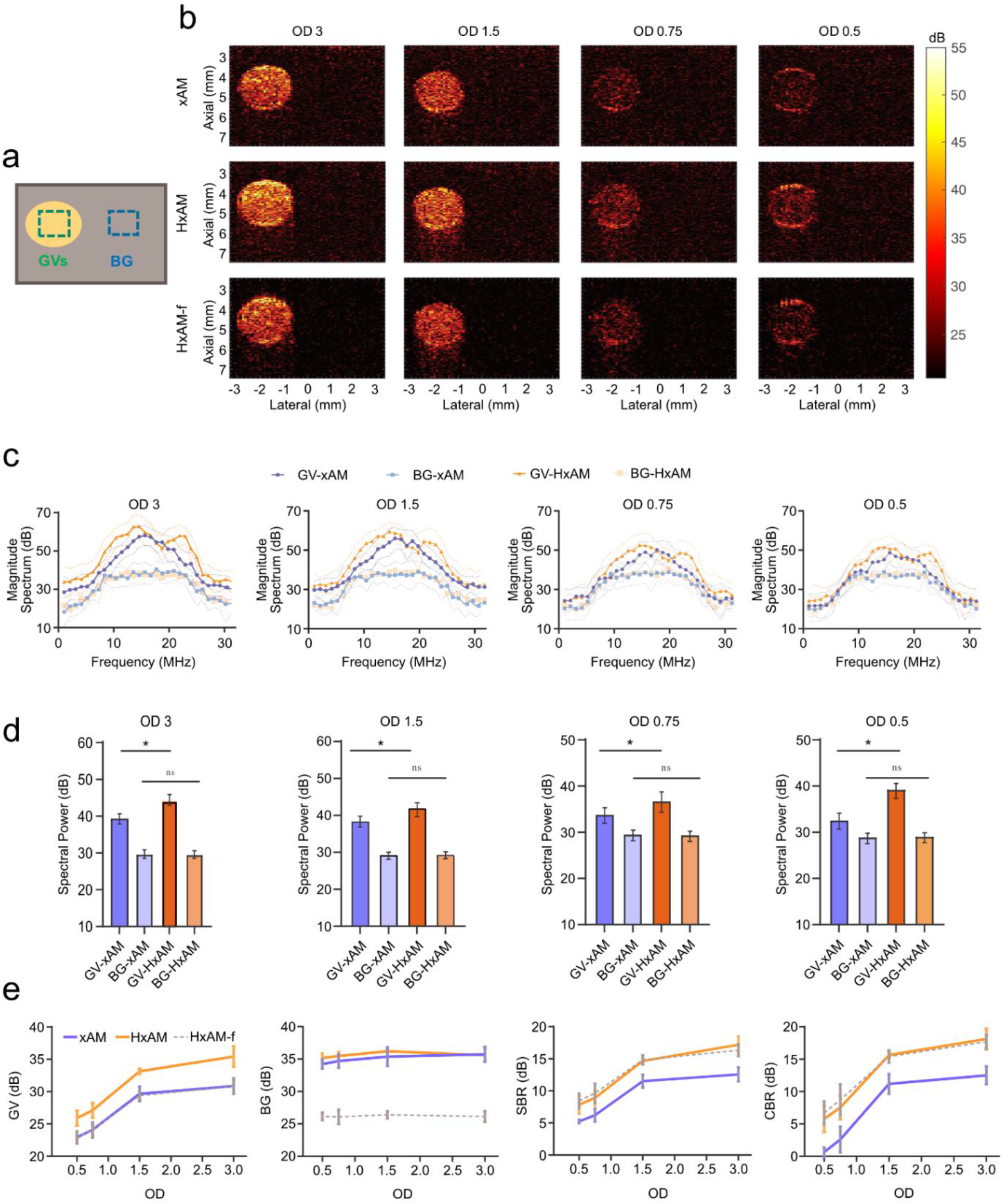
In vitro imaging of GVs at various concentrations in tissue-mimicking phantoms using xAM and HxAM. (a) Schematic of the ultrasound phantom with GV inclusion (yellow) in a tissue-mimicking matrix (gray). ROIs are indicated in green for GVs and blue for the background. The GV inclusions are positioned at a depth of 4.5 mm and have a radius of 1 mm. **(b)** Representative xAM (top row) and HxAM (middle row) images of the same cross-section at different GV concentrations. The bottom row shows HxAM images high-pass filtered at a 17.5 MHz threshold to highlight harmonic contributions. All images are displayed using the same dynamic color range. **(c)** Magnitude Fourier spectra for GV and background signals in xAM and HxAM images at various GV concentrations. **(d)** Quantitative analysis of GV and background signals in xAM and HxAM images as related to GV concentration, including STR and CTR performance metrics. **(e)** Spectral power from the Fourier spectrum of GV and background signals in xAM, HxAM, and HxAM-filtered images at different GV concentrations. Data from N=5 samples; error bars represent the standard error of the mean.

HxAM yielded considerably enhanced images compared to conventional xAM [Fig. 2 (b)]. Spectral analysis revealed a distinct second harmonic signal at 25 MHz, which was solely attributable to the GVs and was not present in the tissue-mimicking background [Fig. 2 (c), (d)]. These attributes translated into a notable increase in signal to background ratio (SBR) and contrast-to-background ratio (CBR), with improvements of 4.7 dB and 5.6 dB, respectively. Further, such enhancements stemmed from an increase in the GV signal, while background levels remained the same between the two methods [Fig. 2 (e)]. HxAM maintained a performance advantage of over 5 dB compared to xAM regardless of averaging parameters. With just 15 averaging repeats, HxAM surpassed 50 such repeats of xAM in imaging quality [Supplementary Fig. 1]. The resulting increase in imaging frame rate can be especially useful for *in vivo* applications and dynamic imaging.

The filtering in HxAM-f substantially suppressed background signal, while also reducing GV signal, resulting in overall SBR and CBR similar to unfiltered HxAM [Fig. 2 (b), (e)]. To confirm that the distinct harmonic signals observed in HxAM can be attributed to the nonlinear buckling behavior of GVs, we also imaged stiff-shelled GVs that do not exhibit robust buckling behavior^11^ due to the presence of GvpC, and saw a lack of second harmonic output [Supplementary Fig. 2].

In contrast to HxAM, harmonic imaging using the parabolic focusing of pAM did not significantly increase SBR or CBR compared to fundamental imaging [Supplementary Fig. 3], consistent with our previous report^16^. This difference arises from a lower harmonic-to-fundamental signal ratio (0.69, compared to 0.98 in xAM), which we attribute to artificially elevated fundamental signal at full transmit amplitude caused by cumulative nonlinear propagation^17^. In contrast, xAM imaging by design has lower propagation nonlinearity, providing a higher harmonic-to-fundamental ratio for harmonic imaging and a more faithful representation of scatterer locations in the medium.

To ensure that the enhanced contrast of HxAM does not arise from differences in focal dimensions, we mapped the beam profiles for xAM and HxAM transmissions using an acoustic hydrophone. The elevational and lateral profiles of xAM and HxAM imaging showed no major differences. The full width half maximum (FWHM) values for the elevational beam profile were 0.866 mm for xAM and 0.855 mm for HxAM, while for the lateral beam profile, they were 0.806 mm and 0.785 mm, respectively [Supplementary Fig. 4]. Although HxAM has slightly more pronounced side lobes, their amplitude remains below the GVs’ buckling threshold at our transmit pressures, minimizing the likelihood of artifact generation.

To evaluate the utility of HxAM imaging in applications of GVs as acoustic reporter genes, we imaged MDA-MB-231 cancer cells genetically engineered to express GVs^5^ [Fig. 3 (a)]. This cell line is commonly used to model breast cancer in vivo. HxAM improved the detection of cells across a wide range of concentrations. At the highest densities – 3×10^6^ and 3×10^7^ cell/ml – HxAM increased SBR and CBR relative to xAM [Fig. 3 (b)-(d)]. At lower densities where individual cells are expected to be separated within the field of view, HxAM facilitated the identification of a larger number of signal sources (putative cells) compared to xAM – identifying 2.4-times to 3.0-times more sources [Fig. 3 (b), (e)]. To ensure that the contrast points contained GVs, we applied ultrasound pulses at a high pressure (3.6 MPa) using focused parabolic delays exceeding the GV’s irreversible collapse threshold (∼570 kPa) and documented a loss of scattering [Supplementary Fig. 5].

**Figure 3:**
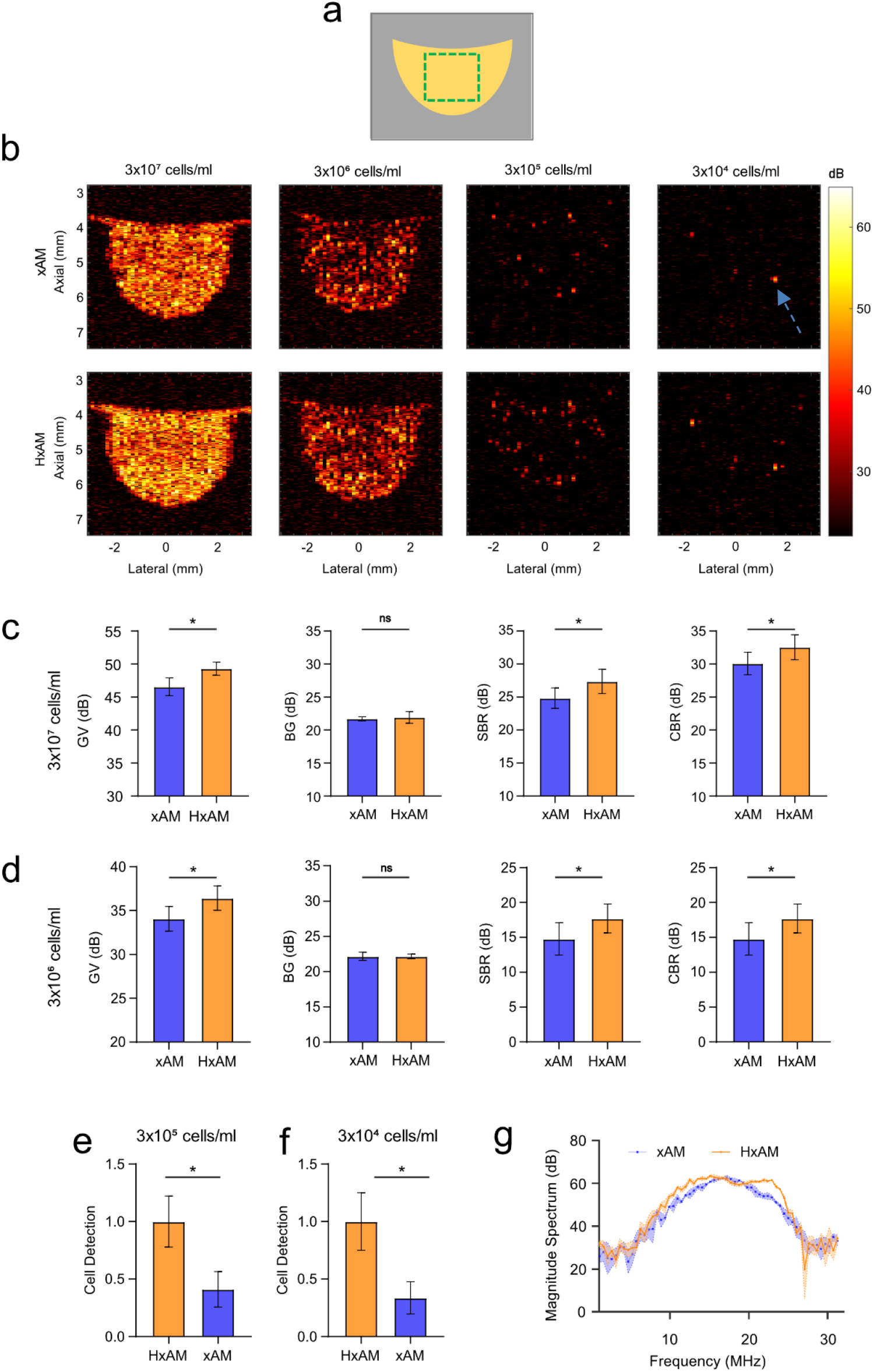
Comparative ultrasound imaging of engineered mammalian cells expressing acoustic reporter genes in vitro. **(a)** Schematic Representation: Mammalian cells (in yellow) embedded within an agar phantom (gray). **(b)** xAM and HxAM Imaging: Images of mammalian cells genetically engineered to express acoustic reporter genes, set in agar at various cell concentrations. All images are displayed using the same dynamic color range. **(c, d)** Signal Analysis: Quantitative evaluation of nonlinear ultrasound signals from xAM and HxAM imaging at concentrations of 3×10^7^ and 3×10^6^ cells, respectively. Analysis includes assessment of gas vesicle (GV) signal from a specified region of interest (ROI) shown in (a) and background signal from the same ROI post-GV acoustic collapse. Bar graphs show GV signals with performance metrics like STR and CTR. **(e, f)** Cell Count Analysis: Cell counts in HxAM and xAM images at concentrations of 3×10^5^ and 3×10^4^ cells, respectively, normalized to HxAM counts at each concentration. Data based on N=5 samples; error bars denote standard error of the mean. **(g)** Fourier Spectral Analysis: Magnitude Fourier spectra derived from xAM and HxAM imaging of a single cell,

To assess the efficacy of HxAM *in vivo*, we imaged mice during intravenous administration of purified GVs, focusing on the liver, where GVs are taken up and degraded as a part of the organ’s phagolysosomal function^2,25^. To evaluate the detection sensitivity, we injected 100 μl solutions containing GVs at either a standard OD of 30 or a minimally detectably concentration of OD 10 [Fig 4 (a)]. Further, to enable imaging of the same exact plane with HxAM and xAM and eliminate confounds from physiological motion, we euthanized the animals immediately before imaging. HxAM allowed GV uptake to be imaged substantially deeper inside liver tissue at both standard [Fig. 4 (b)] and low GV concentrations [Fig. 4 (c), Supplementary Fig. 6 (a)], extending detection depth by up to 20% [Fig. 4 (d)]. The GV-based source of the contrast was confirmed by collapsing the GVs with high-pressure pulses. This improvement was not due to deeper penetration of the transmit pulse; the expected difference in attenuation between 15.625 and 12.5 MHz is only 0.78 dB over 5 mm of liver tissue^26^. Spectral analyses confirmed the presence of harmonic signals in HxAM [Fig. 4 (e), Supplementary Fig. 6 (b), (c)]. HxAM imaging improved SBR and CTR by 4.32 dB and 11.1 dB at OD 30 [Fig. 4 (f)], and 4.4 dB and 15.8 dB at OD 10 [Fig. 4 (g)], relative to xAM.

**Figure 4:**
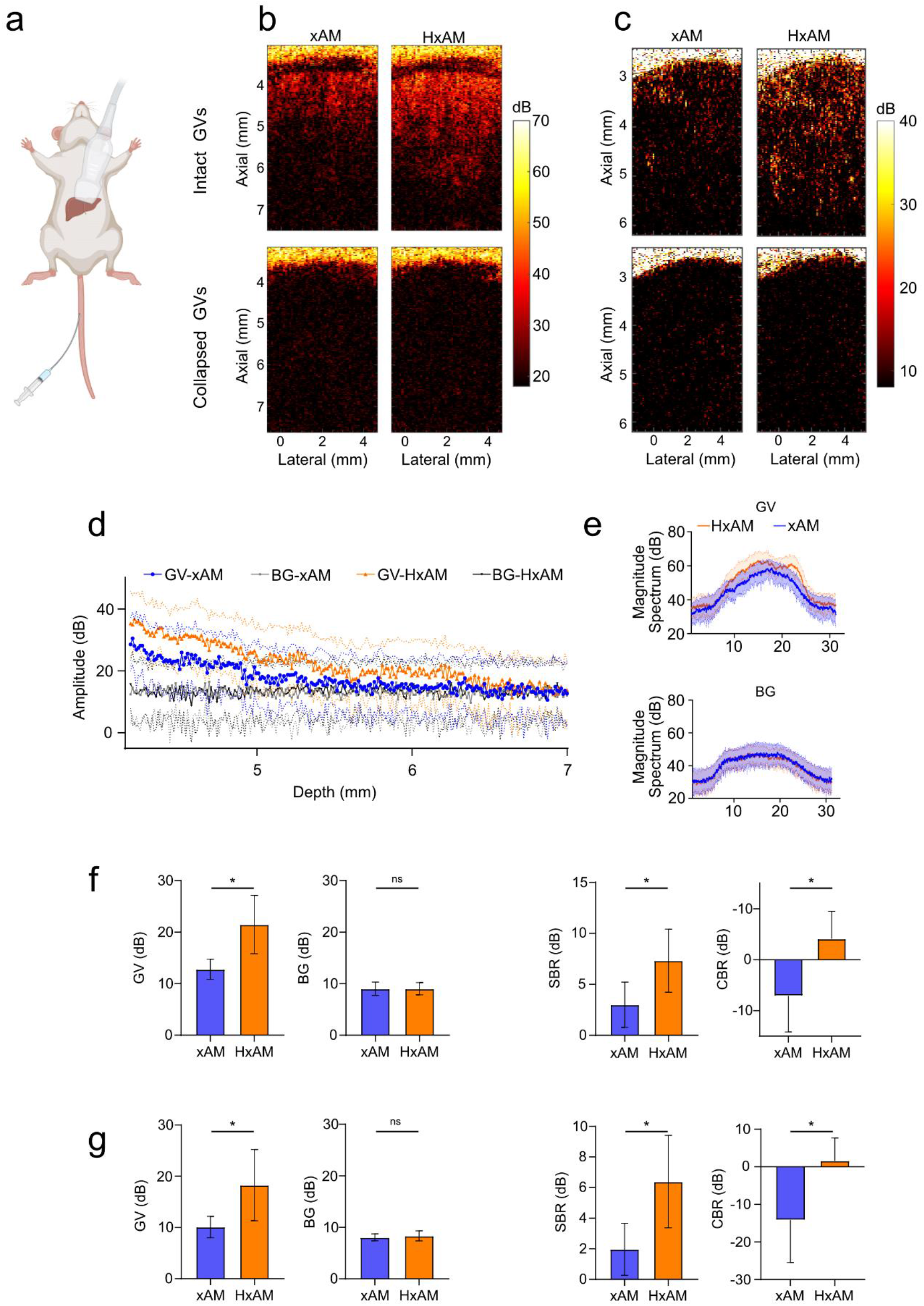
In vivo ultrasound imaging of mice liver after intravenous injection of purified GVs. **(a)** Schematic illustration of the ultrasound mouse liver experiment. **(b, c)** Representative examples of xAM and HxAM images of mice liver, acquired at the same cross-section using both imaging techniques at OD_500_ 30 and 10, respectively. Top and bottom rows correspond to detection of GVs intact and after acoustic-based collapse, **respectively. (d)** Axial line plots corresponding to GV and background signal at varying depths of images in (b). The solid line plots and the corresponding dotted double-sided bands represent mean and the standard error, respectively, estimated across all columns of the ultrasound images in (b). **(e)** Magnitude Fourier spectra associated with GV and background signals in xAM and HxAM images reported in (b). **(f**,**g)** Quantitative analysis of GV and background signal in mice liver using xAM and HxAM imaging, at OD_500_ of 30 and 10, respectively. N=5 mice at each GV concentration and the error bars represent standard error of the mean.

Taken together, the results of this study suggest that HxAM imaging improves the detection of GVs over xAM in all the main *in vitro* and *in vivo* scenarios. Harmonic frequencies are selectively amplified in the GV signal without altering background, leading to marked improvements in SBR and CBR. We anticipate that this enhanced sensitivity will facilitate *in vivo* cell imaging, while its specificity for buckling GVs over stiff GVs will contribute to dynamic biosensing^12,27^. The enhanced performance of HxAM is attributed to the capture of harmonic signals arising from GV scatterers, with minimal background from nonlinear propagation artifacts (unlike pAM).

HxAM imaging inherits some limitations of xAM imaging, including a reduced lateral and axial field of view. In addition, it adds the requirement for transducers and scanners with relatively high transmit-receive bandwidth. However, HxAM’s improved ability to detect GVs and GV-expressing cells makes it an attractive imaging method for biomolecular and cellular ultrasound.

## METHODS

### Ultrasound acquisition sequence

We use a Verasonics Vantage ultrasound scanner, equipped with an L22-14vX probe, to execute various imaging sequences such as xAM, pAM and its respective harmonic versions. This probe features a linear array composed of 128 elements, each spaced at a 0.10-mm interval. It has an elevation focus of 8 mm, a 1.5-mm elevation aperture, and operates at a central frequency of 18.5 MHz, offering a bandwidth of 67% at −6 dB. For transmission, we use single-cycle waveforms at frequencies of 15.625 MHz and 12.5 MHz for each active element in the array, which ensures that our base frequency is divided by factors of 4 and 5 respectively, in sync with the system’s 62.5-MHz sampling rate. For harmonic versions of xAM and pAM, we opt for a transmit frequency of 12.5 MHz, while maintaining a receive sampling rate of 62.5 MHz—the highest rate achievable on the Verasonics Vantage system. To balance lateral view and axial depth, the xAM sequence employs an aperture consisting of 65 elements, centering the array but silencing the middle element to allow for symmetric AM coding. This setup results in 64 ray lines for each xAM image. For the pAM sequence, we adjust the focal depth of the parabolic delays to 4.5 mm, aligning it with the center of the GV cylindrical inclusion. We use a 38-element aperture for this sequence, resulting in an F-number of 1.125 and producing 89 ray lines to optimize the dimensions of the pAM images. The raw radio frequency (rf) data is collected and processed through a custom-built, real-time image reconstruction pipeline, as described in the paper^17^. This includes a specialized beamforming algorithm tailored to meet the unique demands of xAM imaging. Further, the signal to noise ratio (SNR) of the acquired radio frequency (RF) data was enhanced using repeated transmit-receive events, which involves accumulating the acquired data at the same receive location within the dRAM memory to suppress background noise in the acquired signals. We selected 15 averaging repeats as the standard for both HxAM and xAM imaging techniques, as per experiments and analysis performed in Supplementary Figure 1.

Lastly, for both xAM and HxAM imaging techniques, the received RF data corresponding to the three AM transmit sequences were accumulated at the same location in the receive buffer of the scanner. We applied an apodization of -1 to each of the two half-amplitude receives of the AM sequences, aiding in the process of AM cancellation. The reasoning behind aggregating the received data from the three different transmit sequences at a single location, as opposed to collecting them at separate locations and then subtracting them during post-processing, was to ensure there was no saturation of the receive buffer, which has a 14-bit dynamic range. This approach was particularly crucial when working with a higher number of averaging repeats. If the buffer became saturated, especially in the data corresponding to the full amplitude transmit of the AM sequence, it could lead to imbalances in AM cancellation, which in turn would result in considerable image artifacts.

### Preparation of gas vesicles

Gas vesicles were isolated from *Anabaena sp*. as detailed in prior studies^8^. To strip the native GVs, a 6M urea solution was applied, followed by repeated rounds of centrifugation for stripped GVs. The medium was then exchanged through four rounds of flotation and subsequent removal of the subnanat, yielding rounds of dialysis in PBS to guarantee the thorough removal of the native GvpC layer. The concentration of GVs was quantified by measuring the optical density at 500 nm (OD_500_) using a NanoDrop ND-1000 spectrophotometer (Thermo Fisher Scientific, Waltham, MA, USA). For *Anabaena sp*., an OD_500_ of 1 is equivalent to a GV concentration of 184 pM or a volume fraction of 0.04% in an aqueous suspension^8^. This quantification is based on the principle that purified GVs suspended in PBS scatter visible light, allowing their concentration to be accurately measured by assessing the OD at 500 nm.

### In vitro imaging of tissue-mimicking phantoms

Tissue-mimicking phantoms were prepared using a mixture of 1% agarose by weight/volume (w/v) in PBS and 0.2% AlO_3_ (w/v)^16–18^. Specialized 3D-printed molds were utilized to form cylindrical wells with a diameter of 2 mm. GVs were briefly heated at 42°C for one minute and subsequently combined at an equal ratio with low-melt agarose that was also at 42°C. This mixture was then loaded into the phantom wells to achieve various GV concentrations corresponding to optical density values of 3, 1.5, 0.75, and 0.5 at a wavelength of500nm. The AlO_3_ level was specifically selected to emulate the echogenic scatter properties of the GVs, as determined by the contrast-to-noise ratio in B-mode ultrasound imagery. The center of these GV-filled cylindrical wells was positioned at a 4.5mm depth. All these phantoms were imaged while positioned on an acoustic-absorbing material and submerged in PBS. All phantom experiments were conducted across N=5 samples, for different ODs and GV types.

### In vitro imaging of engineered MDA cells expressing acoustic reporter genes

For all *in vitro* cellular imaging experiments, we used MDA-MB-231-mARGAna cells cultured in DMEM supplemented with 10% TET-free FBS and 1X penicillin/streptomycin^5^. For xAM and HxAM imaging of MDA-MB-231-mARGAna cells suspended in agarose phantoms, cells were cultured in T225 flask with 30 mL media. Cells were seeded and induced with 1 µg/mL doxycycline after an overnight incubation and at subsequent days as indicated, except for the uninduced control which was grown in a 10 cm dish without doxycycline. Media was changed daily thereafter until cell harvest. Cells were trypsinized with 6mL 0.25% Trypsin/EDTA for 6 minutes at 37 °C, after which the trypsin was quenched by addition of 8 mL media. Cells were harvested and resuspended at 60,000,000 cells/mL. 10-fold serial dilutions were performed with each cell line. Each cell dilution was mixed 1:1 with 2% low-melt agarose before loading into agarose phantom wells (5 replicates each). Cells were imaged with an L22-14vX transducer at 0.5 MPa for both xAM and HxAM imaging.

### In vivo imaging of mice liver

The *in vivo* experiments were performed on Cg-Foxn1^nu^/J female mice (Jackson Laboratory) aged 7 weeks under a protocol approved by the Institutional Animal Care and Use Committee of the California Institute of Technology. This study did not require any randomization or blinding techniques. The mouse was sedated using a 2%–3% isoflurane anesthesia. The experimental procedures involved administering an intravenous injection of 100 μL of a sterilized GV solution dissolved in PBS^25^. We tested two distinct concentrations of GVs, with OD_500_ of 10 and 30, examining a total of five animals for each GV concentration. After waiting for 25 minutes to allow for liver phagocytosis of the GVs, as supported by previous research, the animal was then humanely euthanized. Subsequently, we obtained xAM and HxAM images right away. Thereafter, we collapsed the GVs using high acoustic transmit pressure (>3.5 MPa), and reacquired xAM and HxAM images at the same liver cross-section. The reason for performing imaging post-euthanasia was to eliminate any physiological motion from breathing or heartbeat. Such motion could introduce variability in the imaging cross-section, thus compromising the reliability of the comparison between xAM and HxAM images.

### Quantitative and statistical analysis

Quantitative numbers to assess performance of the AM imaging techniques were computed using SBR 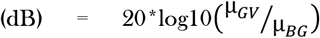,and CBR 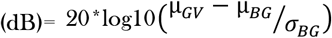, where µ and σ denotes mean and standard deviation, estimated in the identified ROIs. The Fourier spectra for both GV and BG signals were calculated using the same ROIs as those applied for other quality metrics. Columns within each ROI and across various samples were collectively aggregated as repeated measures for estimating the frequency content. The Fourier spectra for these aggregated column data were computed using the ‘fft’ function in MATLAB. These analyses were performed for all *in vitro* and *in vivo* experiments involving tissue mimicking phantoms, engineered cells and mice.

Additionally, due to the sparse distribution of cells at the lower two concentrations depicted in Figure 3, it was impractical to estimate SBR and CBR. Therefore, we quantified pixels exceeding a specific threshold as contributing to cell presence for each imaging method. We confirmed that the threshold value resulted in a zero cell count in post-collapse images. The cell counts obtained via xAM were then normalized against those acquired through HxAM, with these comparative results reported in Figure 3. Finally, to assess the statistical significance of the comparisons between xAM and HxAM imaging, we employed t-tests. Statistical significance was established at a p-value greater than 0.95, denoted by an asterisk (*).

## SUPPLEMENTARY DATA

### Impact of averaging repeats on xAM and HxAM imaging

To assess the effect of averaging these repetitions on the image quality of xAM and HxAM imaging, we performed a focused experiment. We used a tissue-mimicking phantom with GV inclusion at a concentration of OD 3. We varied the number of averaging repeats in a range from 1 to 50, incrementing in steps of 5 for each test run. This testing was carried out using a set of five phantoms to ensure a robust sample size. The results presented in Supplementary Figure 1 reveal a correlation between the number of averaging repeats and the enhanced performance in both xAM and HxAM imaging modalities. However, the quality of the HxAM images consistently exceeded that of the xAM images, regardless of the number of averaging repeats applied. The quantitative plots show that the GV signal [SFig. 1 (a)] with HxAM imaging is at least 5 dB higher than xAM imaging, while no noticeable difference in the background signal [SFig. 1 (b)] was observed, which resulted in higher SBR [SFig. 1 (c)] and CBR [SFig. 1 (d)]. Further, the image quality metrics show that HxAM imaging with 15 averaging repeats achieved superior results compared to xAM imaging with as many as 50 repeats. Drawing from these findings, we selected 15 averaging repeats as the standard for both HxAM and xAM imaging techniques in the subsequent experiments presented in this paper.

### Harmonic imaging using wild type GVs

To validate the critical role of nonlinear behavior in the effectiveness of HxAM imaging, our study included experiments with wild-type gas vesicles (wtGVs) embedded in tissue mimicking phantoms. Specifically, wtGVs naturally lack nonlinear properties due to the presence of the structurally stiffening protein gvpC. As expected, wtGVs did not demonstrate a perceptible GV signal in AM imaging for both xAM and HxAM imaging [SFig. 2 (a)]. Further, HxAM imaging demonstrated no quantifiable advantage over xAM, contrary to what was observed with stripped GVs, when compared at the same OD [SFig. 2 (b)]. This outcome highlights the importance of nonlinear buckling in the enhanced performance of HxAM imaging. Moreover, the magnitude Fourier spectrum analysis of wtGVs revealed no significant amplification at harmonic frequencies [SFig. 2 (c), (d)], further substantiating our findings.

### Harmonics imaging using parabolic AM pulse sequences

To evaluate the feasibility of using harmonic imaging to enhance the performance of parabolic AM (pAM) pulse sequence, we conducted *in vitro* experiments with GVs embedded in a tissue mimicking phantom at OD 3.5. The GVs inclusions were imaged using both pAM and xAM imaging, along with its harmonic counterparts, at the same phantom cross-section [SFig. 3 (a)]. The acquired channel data corresponding to the three transmit AM pulses were stored individually for comparative analysis. Further, for simplified analysis, the quantified GV, BG signals were normalized by respective pAM and xAM signals. Similarly, the SBR and CBR values were normalized by pAM and xAM counterparts, respectively. The quantitative results [SFig. 3 (b)] reveal a notable difference in the harmonic-to-fundamental frequency ratios of the GV signal between pAM and xAM imaging methods. Specifically, the harmonic-to-fundamental frequency ratio for pAM stands at a lower value of approximately 0.69, while xAM registers a higher value of approximately 0.98. This leads to differing SBR, with pAM at 1.03 and xAM at 1.47. Such differences are critical in the context of limited bandwidth of the transducer and the scanner, suggesting that the replacement of fundamental frequencies with harmonics in pAM does not significantly enhance imaging performance. In contrast, in xAM imaging, the harmonic frequencies are on par with the fundamental frequencies in amplitude, which contributes to a significant improvement in image quality when harmonics are incorporated.

The constraint on the harmonic to fundamental frequency ratio observed in pAM is predominantly due to an artificially increased fundamental frequency component due to propagation artifacts commonly observed with approach. The presence of these artifacts can also be observed in SFig. 3 (a). However, xAM imaging is engineered to mitigate this artifact, making harmonic contributions a substantial contribution. Further examination of the absolute value GV signal in pAM and xAM imaging [SFig. 3 (b)], where assuming a similar harmonic response for both imaging techniques, pAM imaging at 15.625 MHz experiences a reduction of the full amplitude receive signal by 60% after AM cancellation, whereas xAM imaging produces a reduction of up to 80%, thereby improving the harmonic-to-fundamental frequency ratio. This substantial difference underscores the distinctive imaging capabilities and benefits that xAM holds over pAM in harmonic imaging applications.

To assess the potential of harmonic imaging in augmenting the efficacy of the parabolic AM (pAM) pulse sequence, we undertook *in vitro* studies with GVs embedded within a tissue-mimicking phantom at an OD_500_ of 3.5. Imaging of the GV inclusions was performed using both pAM and xAM techniques at 15.625 MHz, as well as their harmonic counterparts (HpAM and HxAM) at 12.5 MHz, across the same section of the phantom, as shown in [SFig. 3 (a)]. The harmonic filtered images (HpAM-f and HxAM-f) were estimated by applying a high pass filter with a cut off frequency of 17.5 MHz. The channel data from the three AM transmit pulses were recorded separately for a detailed comparative evaluation. For ease of analysis, the measured GV and background signals were normalized against their corresponding pAM and xAM signals. Similarly, the SBR values were normalized to their respective pAM and xAM benchmarks. The findings, depicted in [SFig. 3 (b)], highlight a significant disparity in the harmonic-to-fundamental frequency ratios of the GV signal between the pAM and xAM methods. Specifically, the ratio for pAM is notably lower, around 0.69, compared to xAM’s higher ratio of approximately 0.98, leading to a variation in SBR values—1.03 for pAM and 1.47 for xAM. This discrepancy underscores the limited utility of substituting fundamental frequencies with harmonics in pAM, given the constrained bandwidth of the transducer and scanner. In contrast, xAM imaging, where harmonic and fundamental frequencies are comparable in amplitude, markedly improves image quality through the inclusion of harmonic frequencies. The observed limitation in the harmonic to fundamental frequency ratio in pAM primarily results from an artificially elevated fundamental frequency component, a consequence of propagation artifacts typical of this method, also observable in [SFig. 3 (a)]. However, xAM imaging is designed to overcome such artifacts, rendering harmonic enhancements significantly more effective. A further analysis of the absolute GV signal values in pAM and xAM [SFig. 3 (c)] reveals that, assuming a comparable harmonic response in both techniques, pAM imaging at 15.625 MHz sees a 60% reduction in the full amplitude receive signal following AM cancellation. In comparison, xAM imaging achieves a reduction of up to 80%, thus favoring an improved harmonic-to-fundamental frequency ratio. This pronounced difference highlights xAM’s superior imaging capabilities and advantages over pAM in the realm of harmonic imaging.

### Elevational and lateral beam profile measurement

We measured the beam profiles for xAM and HxAM imaging using an HNR-0500 needle hydrophone from Onda Corporation. These measurements were performed using an automated two-axis motorized stage for precise control. The hydrophone was securely mounted at the bottom of a water-filled tank, while the L22-14vX ultrasound probe was affixed to the motorized stage. For the imaging sequences, the xAM and HxAM scripts were run in transmit mode. Specifically, we activated the central cluster of 64 elements for executing the full amplitude transmission of the three-pulse xAM sequence. The motorized stage was pre-programmed to conduct a raster scan with increments of 50 μm in the elevational direction and 100 μm in the lateral direction. Data from the hydrophone was captured using a digital oscilloscope operating at a sampling frequency of 9 GHz. After obtaining this data, we identified the planes corresponding to the peak amplitude levels for both the elevational and lateral beam profiles. These selected profiles are illustrated in SFig. 4. The data reveal that the elevational profiles of xAM and HxAM imaging are closely matched [SFig. 4 (a)], with neither method showing a distinct advantage over the other. The full width half max (FWHM) for the elevational beam profile was 0.866 mm and 0.855 mm for xAM and HxAM, respectively. Lateral profile analysis further indicates that the main lobes are consistent between xAM and HxAM [SFig. 4 (b)]. Similarly, the FWHM for the lateral beam profile was 0.806 mm and 0.785 mm for xAM and HxAM, respectively. While HxAM exhibits more pronounced side lobes, their amplitude does not surpass the buckling threshold of the GVs, even at the full transmission amplitude for the pressures employed in this study. Therefore, it is unlikely that these side lobes would result in artifacts within the amplitude-modulated images.

### In vitro imaging of engineered MDA cells expressing acoustic reporter genes

Supplementary Figure 5 displays *in vitro* ultrasound images of engineered MDA cells, post-collapse of the GVs for xAM and HxAM imaging, at different cell concentrations. These post-collapse images are the background counterparts to those depicted in Figure 3, illustrating the effect of GV collapse on the imaging results.

### In vivo imaging of mice liver

Supplementary Figure 6 depicts the axial line plots corresponding to GV and background signal at varying depths of xAM and HxAM images [Fig. 4 (c)], corresponding to IV injection of GVs at the lower concentration. Analysis of the magnitude Fourier spectra associated with GV and background signals in the xAM and HxAM images [Fig. 4 (c)] show a distinct presence of harmonic contribution for HxAM in the presence of GV, which disappears with the collapse of the GVs, as identified as background in [SFig. 6 (b)].

## ACKNOWLEDGMENTS

This research was supported by the National Institutes of Health (grant R01-EB018975 to M.G.S.) and the Chan Zuckerberg Initiative. M.G.S. is a Howard Hughes Medical Institute Investigator.

## AUTHOR CONTRIBUTIONS

R.N. and M.G.S. conceived this study. R.N. developed sequences for data acquisition and performed all experiments.

M.D. assisted with *in vitro* cellular imaging. B.L. assisted with *in vivo* study. Z.J. assisted with *in vitro* phantom studies.

D.M. produced the GVs. R.N. and M.G.S. wrote the first draft of the manuscript. All authors edited and approved the final version of the manuscript.

## DATA AND MATERIALS AVAILABILITY

The data that supports the findings of this study are available from the corresponding author upon reasonable request.

## COMPETING INTERESTS

The authors declare no competing financial interests.

---

**Supplementary Figure 1:**
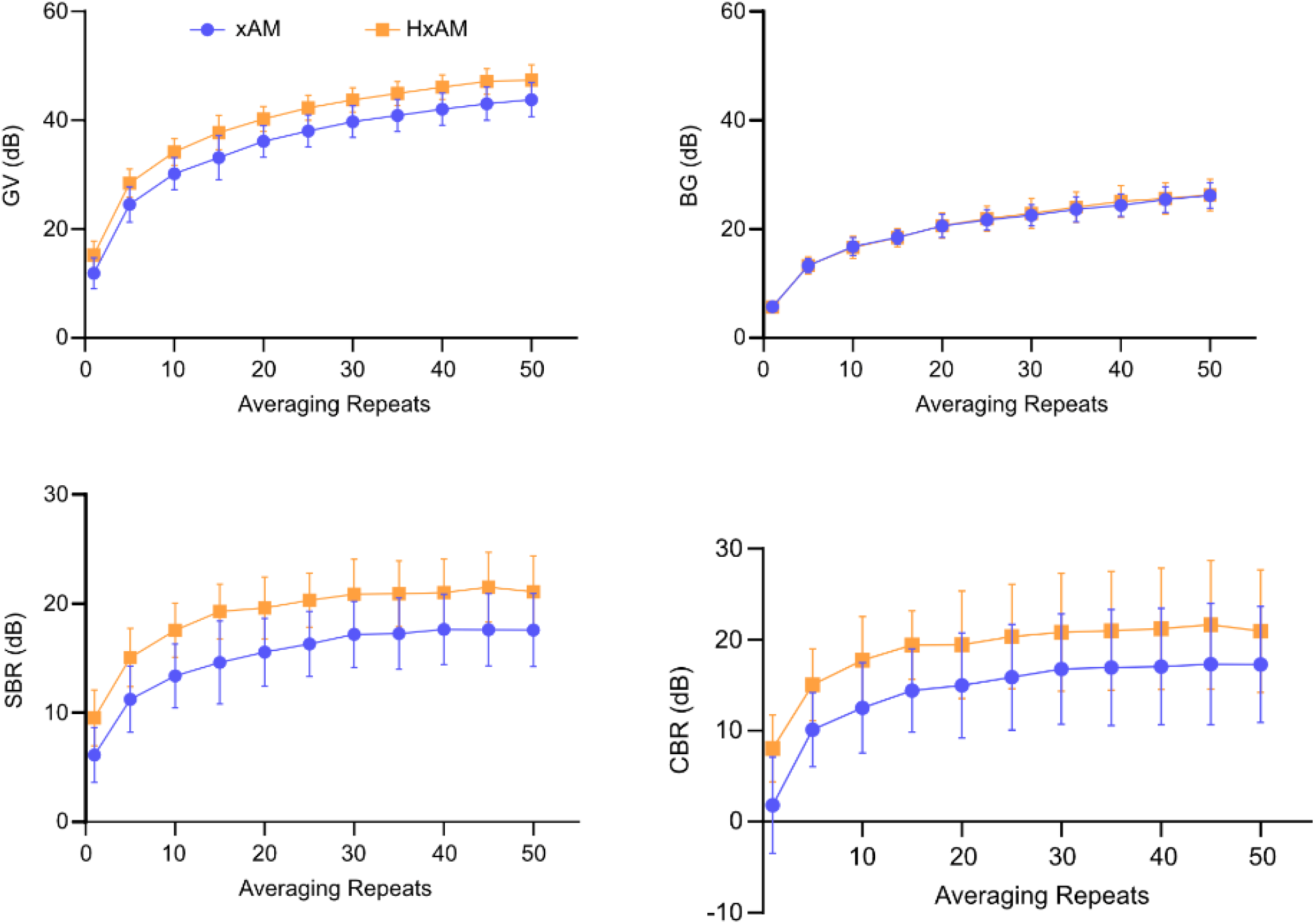
Ultrasound imaging of GV at varying acquisition averaging repeats. The quantitative plots show **(a)** GV signal at OD_500_ 3.5 and the corresponding **(b)** BG signal in the tissue mimicking phantom material as a function of averaging repeats for both HxAM and xAM imaging. The corresponding **(c)** SBR and **(d)** CBR metric. N=5 phantom samples.

**Supplementary Figure 2:**
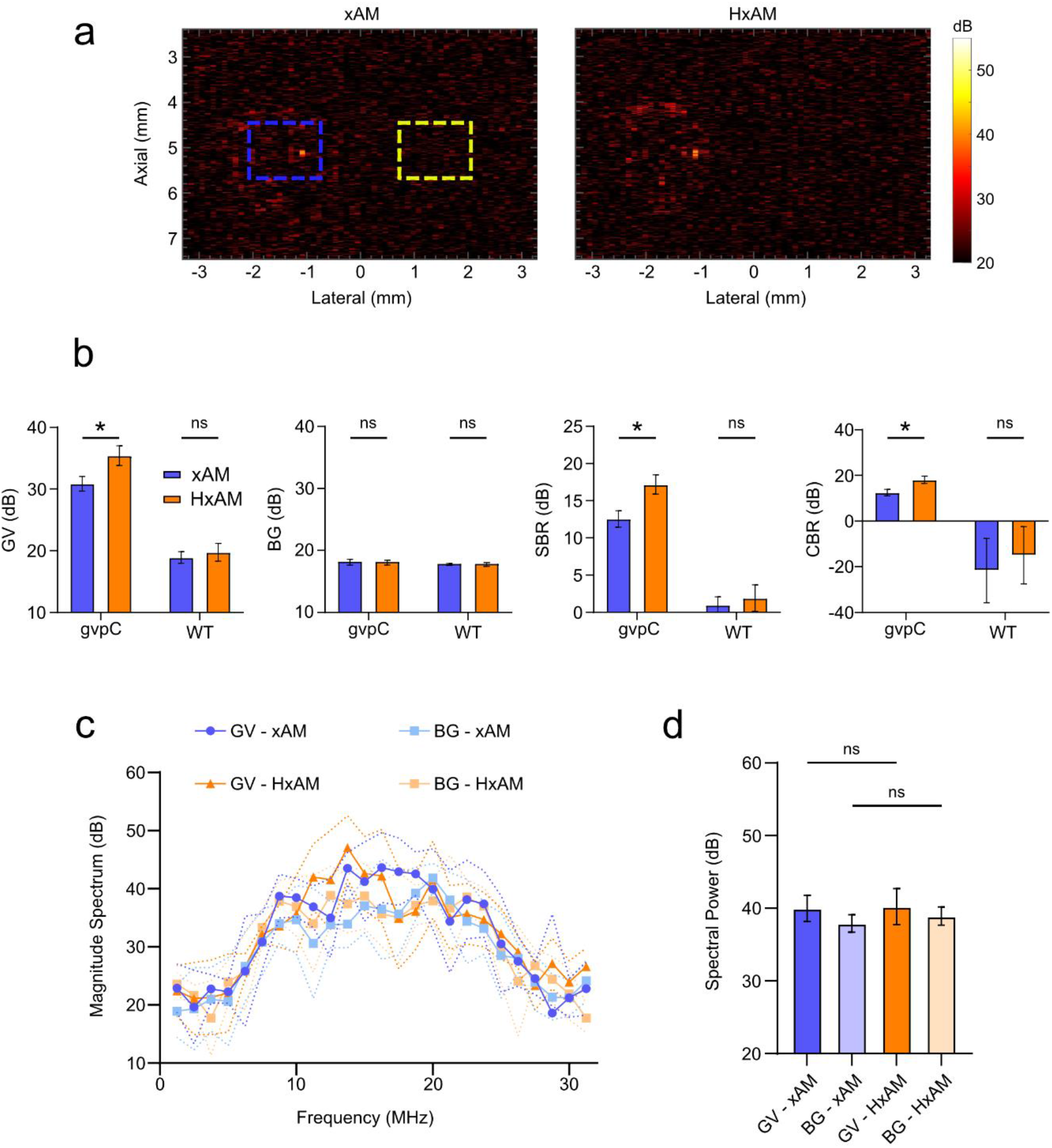
In vitro ultrasound imaging of wtGVs in tissue-mimicking phantoms. **(a)** xAM and HxAM images of wtGVs at a relatively high concentration of OD3. **(b)** Quantitative analysis of xAM and HxAM imaging with respect to wtGVs, corresponding background and comparison with GVs at matched concentration. The bar plots report the magnitude of GV and background signal, and the corresponding performance metrics of STR and CTR. **(c)** Magnitude Fourier spectrum associated with xAM and HxAM images of wtGVs relative to the background, and **(d)** the corresponding spectral power. N=5. Errorbars: standard error of the mean (SEM).

**Supplementary Figure 3:**
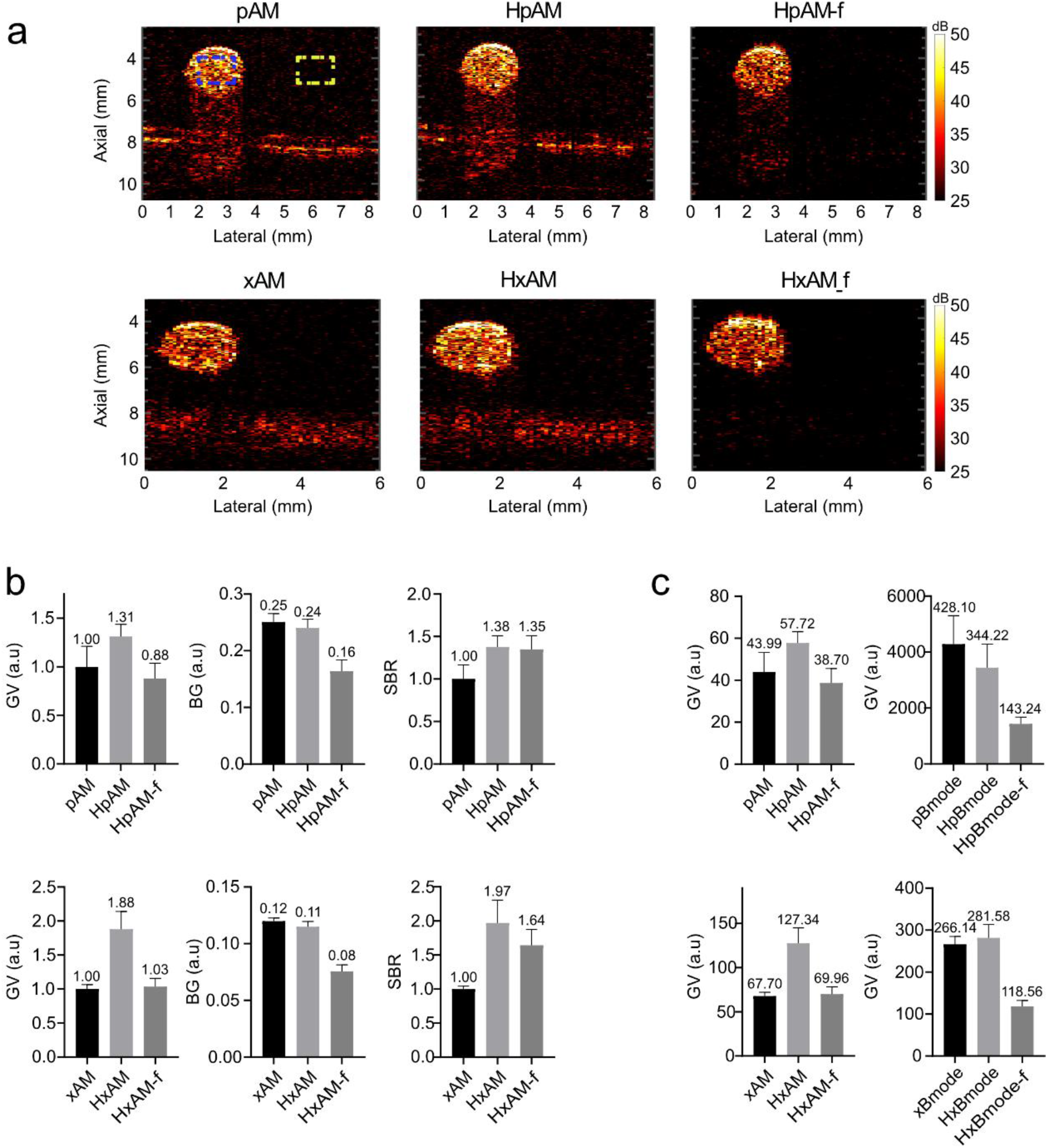
Comparative imaging and signal analysis using a tissue mimicking phantom embedded with a GV inclusion. **(a)** The top row displays images from pAM, HpAM, and harmonic filtered HpAM (HpAM-f) imaging of a GV inclusion at OD 3.5. The bottom row presents the corresponding images for xAM, HxAM, and harmonic filtered HxAM (HxAM-f), captured at the same imaging cross-section. **(b)** Barplots display a quantitative assessment of the GV and background signals within the ROIs identified in (a), along with estimates of the SBR. The top and bottom rows detail the GV and background signals normalized to the GV signal obtained from pAM and xAM imaging at 15.625 MHz, respectively. The SBR values are accordingly normalized against the pAM and xAM counterparts at 15.625 MHz. **(c)** displays the absolute values of the GV signal from (b), juxtaposed with the GV signal captured at full amplitude transmit before AM cancellation. N = 5 samples; error bars represent standard error of the mean.

**Supplementary Figure 4.**
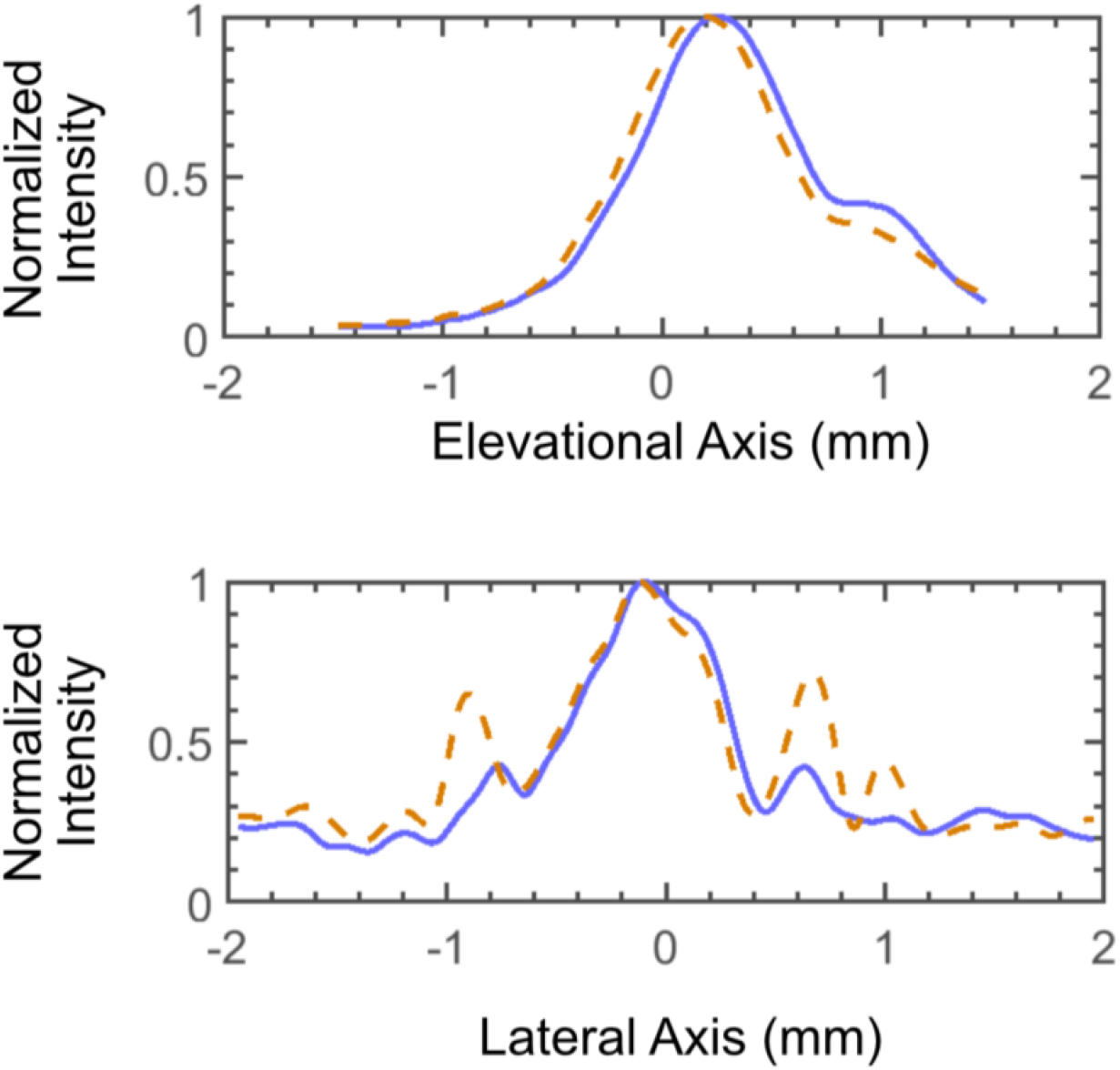
Hydrophone measured beam profile of xAM and HxAM imaging. The top and bottom rows correspond to elevational and lateral beam profiles of xAM (blue) and HxAM (orange) imaging.

**Supplementary Figure 5:**
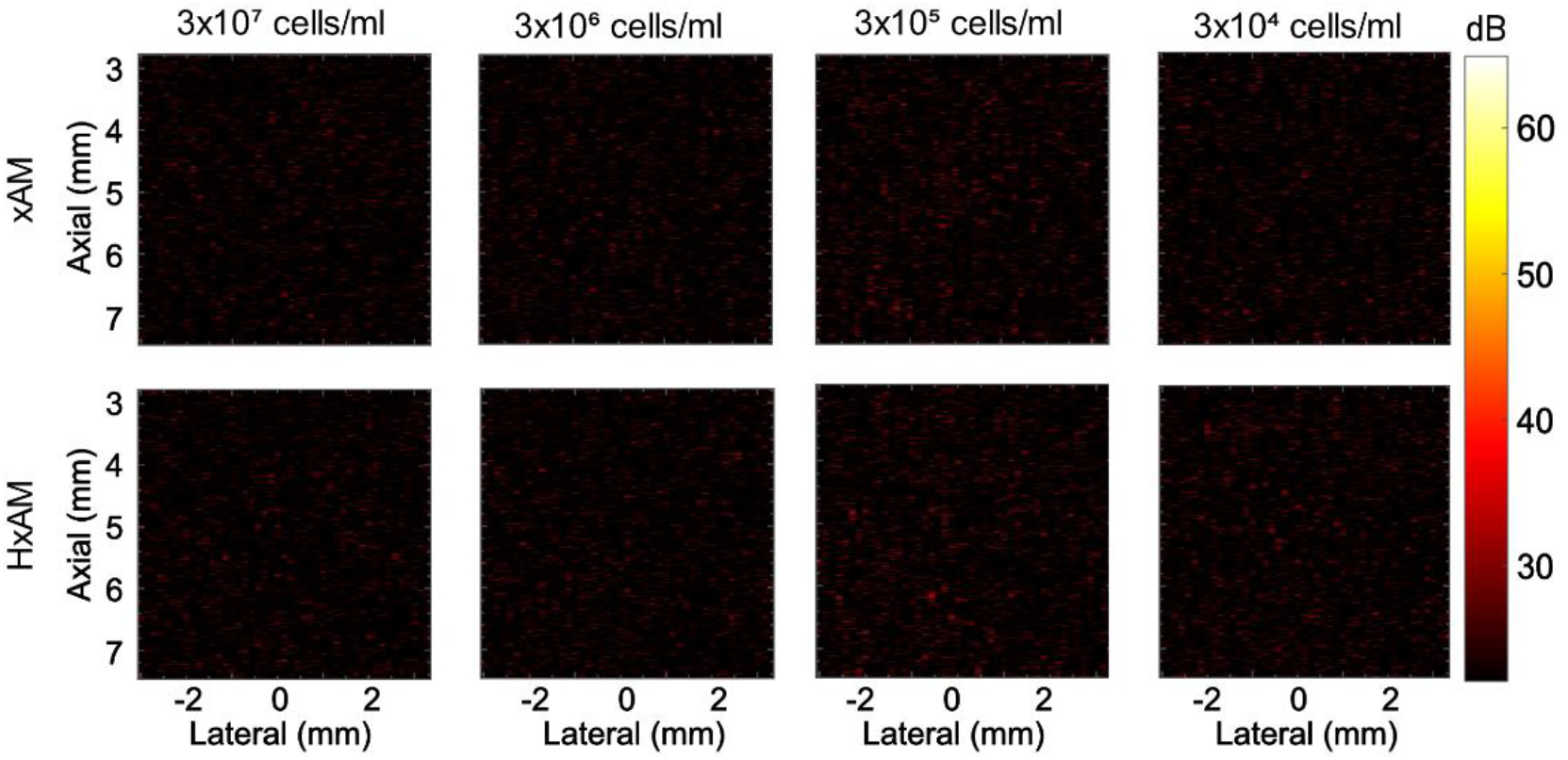
In vitro ultrasound imaging of engineered mammalian cells with collapsed gas vesicles. This figure displays ultrasound images of engineered mammalian cells expressing acoustic reporter genes, following the collapse of the GVs. The top row presents xAM images, and the bottom row shows HxAM images, across various cell concentrations. These images are the post-collapse counterparts to those depicted in Figure 3, illustrating the effect of GV collapse on the imaging results. Data were collected from N=5 sample sets.

**Supplementary Figure 6:**
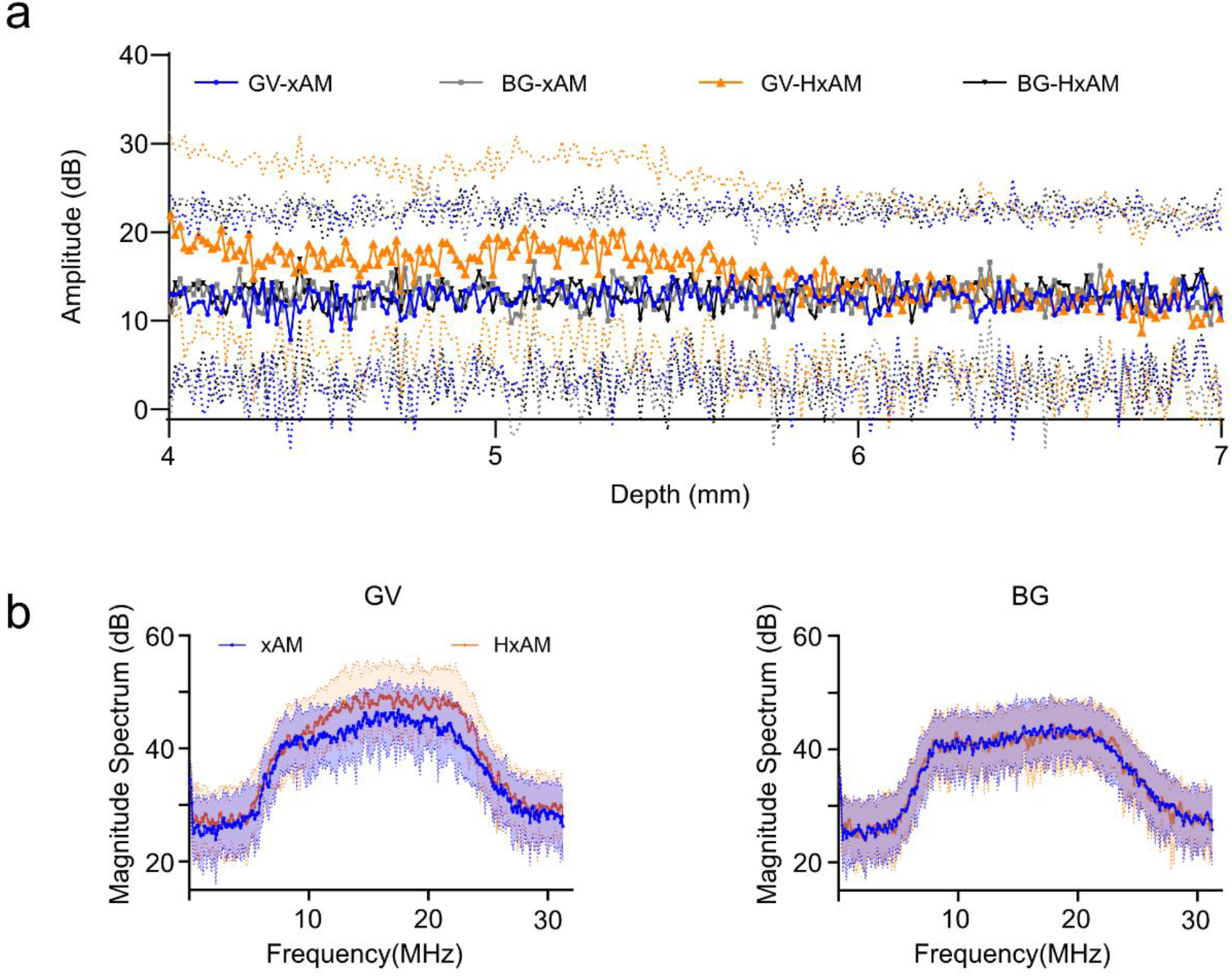
In vivo ultrasound imaging of mice liver after intravenous injection of purified GVs at OD_500_ 10. **(a)** Axial line plots corresponding to GV and background signal at varying depths of images corresponding to IV injection of GVs. The solid line plots and the corresponding dotted double-sided bands represent mean and the standard error, respectively, estimated across all columns of the ultrasound images in Figure 4 (c). **(b)** Magnitude Fourier spectra associated with GV and background signals in xAM and HxAM images reported in Figure 4 (c). N=5 mice and the error bars represent standard error of the mean.

## Notes

### Competing Interest Statement

The authors have declared no competing interest.

## REFERENCES

1 P. Frinking, T. Segers, Y. Luan, and F. Tranquart, “Three Decades of Ultrasound Contrast Agents: A Review of the Past, Present and Future Improvements,” Ultrasound Med. Biol. 46(4), 892–908 (2020).

2 M.G. Shapiro, P.W. Goodwill, A. Neogy, M. Yin, F.S. Foster, D.V. Schaffer, and S.M. Conolly, “Biogenic gas nanostructures as ultrasonic molecular reporters,” Nat. Nanotechnol. 9(4), 311–316 (2014).

3 R.W. Bourdeau, A. Lee-Gosselin, A. Lakshmanan, A. Farhadi, S.R. Kumar, S.P. Nety, and M.G. Shapiro, “Acoustic reporter genes for noninvasive imaging of microorganisms in mammalian hosts,” Nature 553(7686), 86–90 (2018).

4 A. Farhadi, G.H. Ho, D.P. Sawyer, R.W. Bourdeau, and M.G. Shapiro, “Ultrasound imaging of gene expression in mammalian cells,” Science 365(6460), 1469–1475 (2019).

5 R.C. Hurt, M.T. Buss, M. Duan, K. Wong, M.Y. You, D.P. Sawyer, M.B. Swift, P. Dutka, P. Barturen-Larrea, D.R. Mittelstein, Z. Jin, M.H. Abedi, A. Farhadi, R. Deshpande, and M.G. Shapiro, “Genomically mined acoustic reporter genes for real-time in vivo monitoring of tumors and tumor-homing bacteria,” Nat. Biotechnol. 41(7), 919–931 (2023).

6 A.E. Walsby, “Gas vesicles,” Microbiol. Rev. 58(1), 94–144 (1994).

7 F. Pfeifer, “Distribution, formation and regulation of gas vesicles,” Nat. Rev. Microbiol. 10(10), 705–715 (2012).

8 A. Lakshmanan, G.J. Lu, A. Farhadi, S.P. Nety, M. Kunth, A. Lee-Gosselin, D. Maresca, R.W. Bourdeau, M. Yin, J. Yan, C. Witte, D. Malounda, F.S. Foster, L. Schröder, and M.G. Shapiro, “Preparation of biogenic gas vesicle nanostructures for use as contrast agents for ultrasound and MRI,” Nat. Protoc. 12(10), 2050–2080 (2017).

9 P. Dutka, D. Malounda, L.A. Metskas, S. Chen, R.C. Hurt, G.J. Lu, G.J. Jensen, and M.G. Shapiro, “Measuring gas vesicle dimensions by electron microscopy,” Protein Sci. 30(5), 1081–1086 (2021).

10 P.K. Hayes, B. Buchholz, and A.E. Walsby, “Gas vesicles are strengthened by the outer-surface protein, GvpC,” Arch. Microbiol. 157(3), 229–234 (1992).

11 A. Lakshmanan, A. Farhadi, S.P. Nety, A. Lee-Gosselin, R.W. Bourdeau, D. Maresca, and M.G. Shapiro, “Molecular Engineering of Acoustic Protein Nanostructures,” ACS Nano 10(8), 7314–7322 (2016).

12 Z. Jin, A. Lakshmanan, R. Zhang, T.A. Tran, C. Rabut, P. Dutka, M. Duan, R.C. Hurt, D. Malounda, Y. Yao, and M.G. Shapiro, Ultrasonic Reporters of Calcium for Deep Tissue Imaging of Cellular Signals (Bioengineering, 2023).

13 Q. Shen, Z. Li, M.D. Meyer, M.T. De Guzman, J.C. Lim, R.R. Bouchard, and G.J. Lu, 50-Nm Gas-Filled Protein Nanostructures to Enable the Access of Lymphatic Cells by Ultrasound Technologies (Synthetic Biology, 2023).

14 L. Song, X. Hou, K.F. Wong, Y. Yang, Z. Qiu, Y. Wu, S. Hou, C. Fei, J. Guo, and L. Sun, “Gas-filled protein nanostructures as cavitation nuclei for molecule-specific sonodynamic therapy,” Acta Biomater. 136, 533–545 (2021).

15 Y. Hao, Z. Li, J. Luo, L. Li, and F. Yan, “Ultrasound Molecular Imaging of Epithelial Mesenchymal Transition for Evaluating Tumor Metastatic Potential via Targeted Biosynthetic Gas Vesicles,” Small 19(21), 2207940 (2023).

16 D. Maresca, A. Lakshmanan, A. Lee-Gosselin, J.M. Melis, Y.-L. Ni, R.W. Bourdeau, D.M. Kochmann, and M.G. Shapiro, “Nonlinear ultrasound imaging of nanoscale acoustic biomolecules,” Appl. Phys. Lett. 110(7), 073704 (2017).

17 D. Maresca, D.P. Sawyer, G. Renaud, A. Lee-Gosselin, and M.G. Shapiro, “Nonlinear X-Wave Ultrasound Imaging of Acoustic Biomolecules,” Phys. Rev. X 8(4), 041002 (2018).

18 C. Rabut, D. Wu, B. Ling, Z. Jin, D. Malounda, and M.G. Shapiro, “Ultrafast amplitude modulation for molecular and hemodynamic ultrasound imaging,” Appl. Phys. Lett. 118(24), 244102 (2021).

19 D.P. Sawyer, A. Bar-Zion, A. Farhadi, S. Shivaei, B. Ling, A. Lee-Gosselin, and M.G. Shapiro, “Ultrasensitive ultrasound imaging of gene expression with signal unmixing,” Nat. Methods 18(8), 945–952 (2021).

20 F. Tranquart, N. Grenier, V. Eder, and L. Pourcelot, “Clinical use of ultrasound tissue harmonic imaging,” Ultrasound Med. Biol. 25(6), 889–894 (1999).

21 A. Anvari, F. Forsberg, and A.E. Samir, “A Primer on the Physical Principles of Tissue Harmonic Imaging,” RadioGraphics 35(7), 1955–1964 (2015).

22 N. De Jong, A. Bouakaz, and F.J. Ten Cate, “Contrast harmonic imaging,” Ultrasonics 40(1–8), 567–573 (2002).

23 W.K. Chong, V. Papadopoulou, and P.A. Dayton, “Imaging with ultrasound contrast agents: current status and future,” Abdom. Radiol. 43(4), 762–772 (2018).

24 E. Cherin, J.M. Melis, R.W. Bourdeau, M. Yin, D.M. Kochmann, F.S. Foster, and M.G. Shapiro, “Acoustic Behavior of Halobacterium salinarum Gas Vesicles in the High-Frequency Range: Experiments and Modeling,” Ultrasound Med. Biol. 43(5), 1016–1030 (2017).

25 B. Ling, J. Lee, D. Maresca, A. Lee-Gosselin, D. Malounda, M.B. Swift, and M.G. Shapiro, “Biomolecular Ultrasound Imaging of Phagolysosomal Function,” ACS Nano 14(9), 12210–12221 (2020).

26 G. Ferraioli, V. Kumar, A. Ozturk, K. Nam, C.L. De Korte, and R.G. Barr, “US Attenuation for Liver Fat Quantification: An AIUM-RSNA QIBA Pulse-Echo Quantitative Ultrasound Initiative,” Radiology 302(3), 495–506 (2022).

27 A. Lakshmanan, Z. Jin, S.P. Nety, D.P. Sawyer, A. Lee-Gosselin, D. Malounda, M.B. Swift, D. Maresca, and M.G. Shapiro, “Acoustic biosensors for ultrasound imaging of enzyme activity,” Nat. Chem. Biol. 16(9), 988–996 (2020).

